# Fiber Formulation-Dependent Modulation of Gut Microbial Metabolism in Parkinson’s Disease

**DOI:** 10.64898/2026.04.13.718214

**Authors:** Olivia Adele Todd, Lam Dai Vu, Ingmar Aeneas Jan van Hengel, Beth Statkus, Pieter Van den Abbeele

**Affiliations:** Sorridi Therapeutics, Northbrook, IL, USA; Cryptobiotix SA, Ghent, Belgium

**Keywords:** Fiber, prebiotic, gut-brain axis, Parkinson’s Disease, NeuroFiber, constipation, intestinal inflammation, short chain fatty acids, fermentation

## Abstract

Parkinson’s disease (PD) is associated with altered gut-brain signaling, including microbial dysbiosis, intestinal inflammation, and reduced short-chain fatty acid (SCFA) production. Because dietary fibers are selectively fermented by intestinal microbes to generate SCFAs, fiber formulations tailored to the altered intestinal environment in PD offer a strategy to modulate microbial dysfunction. Here, we used the *ex vivo* Systemic Intestinal Fermentation Research (SIFR®) technology platform, which enables assessment of gut microbiome modulation and host-relevant readouts with demonstrated translational relevance, to assess how fiber substrates influence microbial composition and metabolism of fecal microbiota from individuals with PD (n = 6). Fecal samples were incubated for 24 h with single-fiber, multi-fiber, and food-based formulations. Fermentation outputs, including pH, gas, and SCFAs, were quantified, and select formulations were further characterized by profiling microbial community structure and metabolite output. Relative to a parallel untreated control and osmotic laxative comparator, multi-fiber formulations increased SCFA production (∼2-fold, p = 0.001). These effects were accompanied by increased microbial biomass (∼1.5-fold, p = 0.0007), enrichment of fiber-responsive taxa, and coordinated shifts in metabolites associated with gut-brain signaling. Collectively, these findings show that fiber blend complexity and formulation context shape microbial metabolic engagement, supporting formulation-dependent modulation of gut-derived metabolites linked to gut-brain signaling in PD.

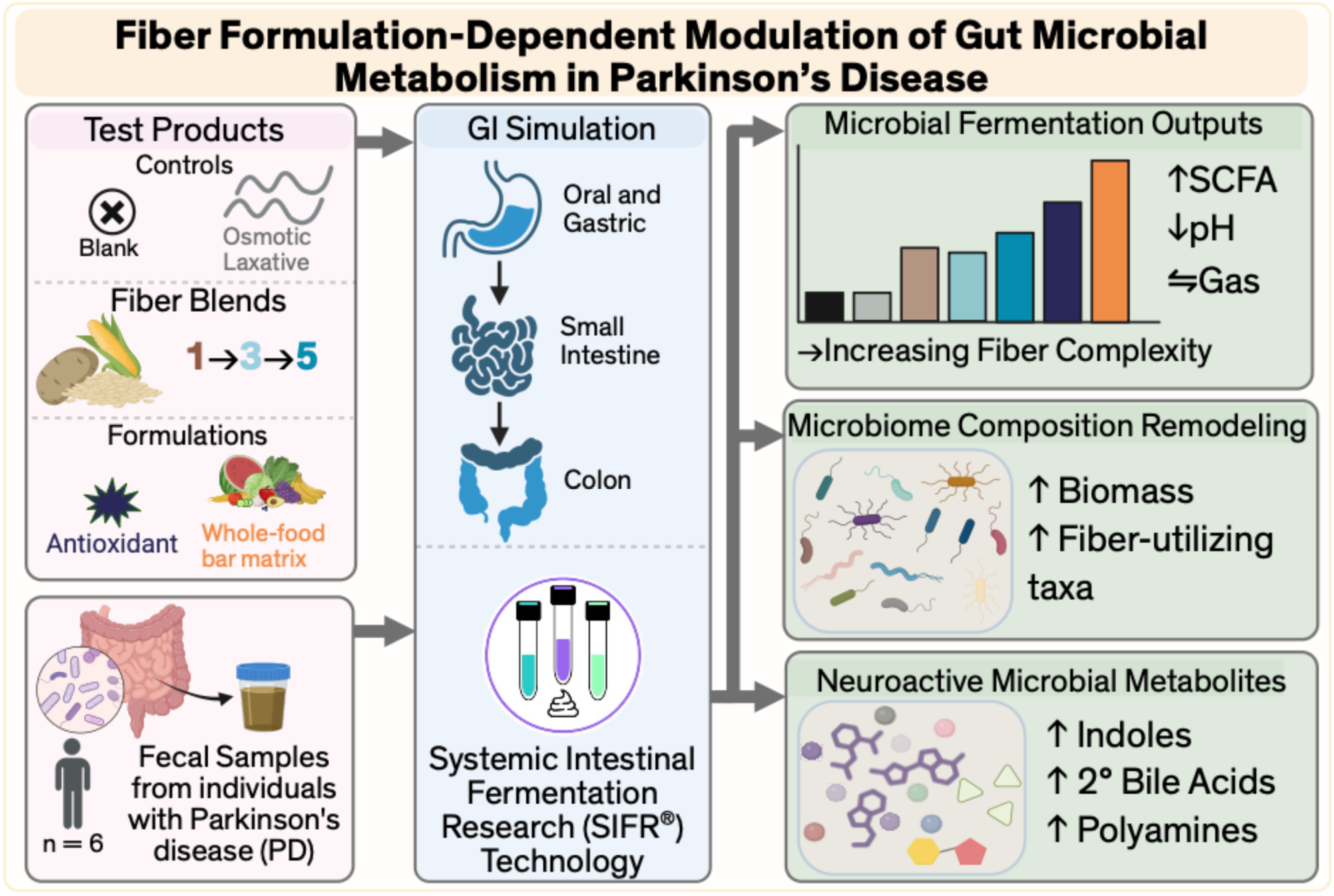

## Introduction

Parkinson’s disease (PD) affects more than 10 million people worldwide, and the global burden of PD is expected to increase substantially with aging populations, with some projections exceeding 25 million cases by 2050.^1^ While PD is primarily thought of as a movement disorder resulting from dopaminergic neuron degeneration, it is increasingly recognized as a complex, heterogeneous condition with widespread non-motor manifestations.^2^ Non-motor symptoms, including sleep dysfunction, mood disturbances, autonomic dysregulation, and gastrointestinal dysfunction, frequently precede motor signs and substantially impact quality of life.^3–6^ Among the non-motor systems affected in PD, the gastrointestinal tract has emerged as a particularly early and biologically informative site of disease involvement.^7,8^

The gastrointestinal tract is intricately connected to the central nervous system through bidirectional neural, immune, endocrine, and metabolic pathways, collectively referred to as the gut-brain axis.^9–11^ The enteric nervous system (ENS) regulates motility, secretion, and local immunity, and is potentially an early site of neuropathology in PD.^12^ PD patients exhibit alterations in gut motility, intestinal barrier integrity, and enteric α-synuclein pathology, suggesting that gastrointestinal dysfunction is not merely symptomatic but may be integral to disease biology.^13–15^ Additionally, microbial communities resident in the gut produce metabolites such as short-chain fatty acids (SCFAs) that influence mucosal immune tone, epithelial health, and neural signaling.^16^ Disruptions in microbial composition and function have been repeatedly observed in PD, with reductions in SCFA-producing taxa and increases in pro-inflammatory organisms reported across multiple cohorts.^17–22^

These microbial and enteric abnormalities are reflected clinically in the high prevalence of gastrointestinal symptoms in PD.^7,23^ Constipation is among the most frequently reported gastrointestinal manifestations of PD, affecting up to 50-80% of individuals, and can precede motor symptoms by years or decades.^24,25^ Prospective cohort studies have demonstrated that reduced bowel movement frequency is associated with increased risk of future PD diagnosis, making constipation not only a clinical burden but a prodromal marker.^7^ Chronic constipation in PD is associated with straining, bloating, and a reliance on laxatives, significantly reducing patient quality of life and increasing caregiver burden.^26^ Accumulating evidence suggests that gastrointestinal dysfunction in PD may occur in the context of low-grade intestinal inflammation. Elevated levels of fecal calprotectin, a neutrophil-derived protein widely used as a non-invasive biomarker of intestinal inflammation^27^, have been reported in individuals with PD compared to age-matched controls.^13,14^ Increased fecal calprotectin reflects mucosal immune activation and has been associated with alterations in intestinal permeability and microbial composition. Notably, intestinal inflammation itself can impair neuromuscular function within the enteric nervous system, disrupt epithelial barrier integrity, and alter secretion and motility patterns, all of which may contribute to delayed colonic transit and constipation.^25^

Clinical management of constipation in PD remains largely symptomatic. Osmotic laxatives, such as polyethylene glycol (PEG), are widely recommended to improve bowel frequency by increasing luminal water content and osmotic load.^28^ However, because these agents do not provide fermentable substrate, they are not expected to engage gut microbial metabolism, support SCFA production, or modulate intestinal immune homeostasis. Prebiotic dietary fibers represent an alternative strategy that may alleviate constipation while supporting gut microbiome function.^29–31^ Fermentation of prebiotic fibers by specific bacterial taxa produces SCFAs, including acetate, propionate, and butyrate, which regulate epithelial barrier integrity, mucosal immunity, host metabolism, and neural signaling through immune, endocrine, and direct neuronal pathways.^16,30,32^ Consistent with these mechanisms, both human and animal studies demonstrate that increased dietary fiber intake enriches SCFA-producing taxa, elevates fecal and circulating SCFA levels, and is associated with improved bowel motility and reduced intestinal inflammation.^33–35^ However, the biological effects of prebiotic fiber depend strongly on fiber type and combination, and typical Western diets provide substantially less fermentable substrate than recommended for optimal microbial metabolism. Current dietary guidelines for the USA suggests adults should consume between 22-34 g of fiber per day,^36^ but the average American only consumes ∼17 g of fiber a day.^37^ Therefore, targeted supplementation with microbiome-active fiber formulations may represent a practical strategy to compensate for insufficient dietary fiber intake.

In this study, we used an *ex vivo*, bioreactor-based technology platform (SIFR^®^ Technology) inoculated with fecal microbiota from individuals with Parkinson’s disease to examine how microbiome-active substrates shape microbial metabolism and community structure, using a commonly prescribed osmotic laxative as a non-fermentable clinical comparator. We performed a functional screen across multiple formulations, quantifying pH, gas production, SCFAs, and branched-chain fatty acids (BCFAs), and selected a subset for deeper characterization using microbiome sequencing and untargeted metabolomics. Our objective was to define formulation-dependent effects on microbial ecology and metabolite output relevant to intestinal homeostasis and gut-brain axis signaling in PD.

## Materials and Methods

### Donor Selection and Fecal Sample Collection

Fecal samples were collected under approval from the University Hospital Ghent Ethics Committee (reference number BC-09977). Six donors were selected based on the following criteria: diagnosis of Parkinson’s Disease, between ages 50-75 years old, no antibiotic use for preceding 3 months prior to sampling, no reported probiotic use, non-smoking, low alcohol consumption (<3 units/day), BMI under 30, no known gastrointestinal diseases or disorders. Donor demographics are detailed in **Supplemental Table S1.** Fecal samples were inoculated directly after collection with no cryopreservation or freezing step.

### Gut Inflammation and Barrier Integrity Testing

Fecal calprotectin was measured from donor stool using the fCAL® ELISA (K6927, Bühlmann, Schönenbuch, Switzerland) following the manufacturer’s instructions.^38,39^ Inflammation levels were determined based on established cutoff values: <50 μg/g, none/healthy; 50-120 μg/g, mild; 120-200 μg/g, moderate; >200 μg/g, severe.^14,27,40^ Fecal zonulin was measured from donor stool using IDK® ELISA (K5600, Immundiagnostik AG, Bensheim, Germany) following the manufacturer’s instructions.^38^

### Test Products and Controls

Fermentation substrates and their corresponding human equivalent doses (g/day) are listed in **Table 1**. As comparator interventions, an osmotic laxative (lax), polyethylene glycol 3350 (PEG 3350; MiraLAX^®^, Bayer U.S.) and Psyllium Husk (PH) (Metamucil 4-in1 sugar-free powder, Proctor & Gamble, USA) were included to represent commonly used non-fermentable laxative and single-fiber approaches, respectively. To evaluate how substrate complexity influences microbial responses, we tested two multi-fiber blends (3-Fiber Blend, B3; 5-Fiber Blend, B5). The 5-Fiber Blend was further formulated with additional components, including an antioxidant (5-Fiber + Antioxidant, B5+AO) or within a whole-food bar matrix (5-Fiber + Bar, B5+Bar), to assess how formulation context alters microbial fermentation and metabolite output. For the bar formulation, the amount added was adjusted so that the delivered dose of the 5-Fiber Blend matched that of the powder formulation, with the remaining mass representing the whole-food excipients. Full ingredient compositions are provided in **Supplemental Table S2**.

**Table 1:**
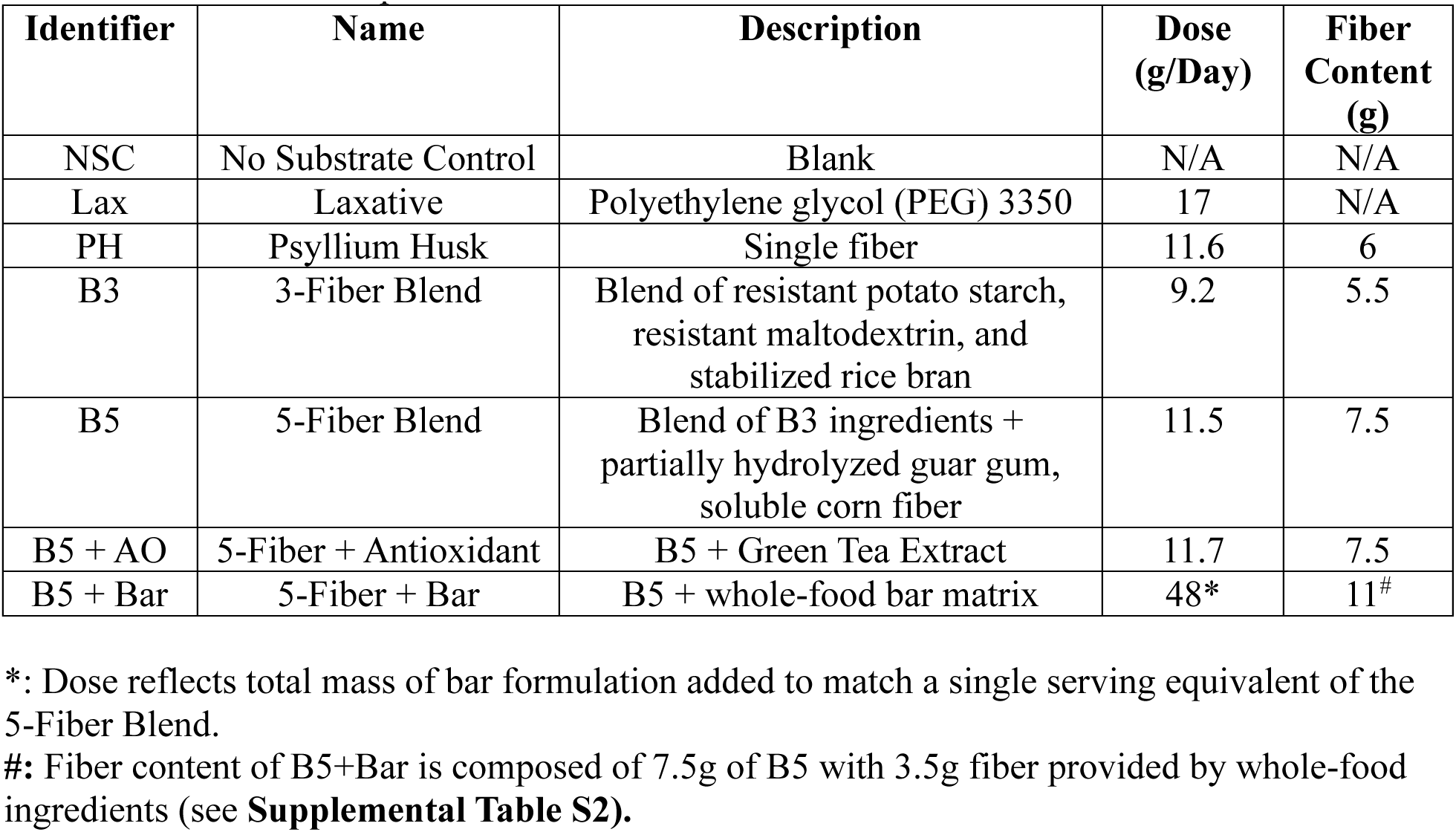
Description of test products used as fermentation substrates, dosage used, and relative amount of fiber (g) per dose. Dosages were determined based on the manufacturer’s serving size recommendation of each product.

| Identifier | Name | Description | Dose (g/Day) | Fiber Content (g) |
| --- | --- | --- | --- | --- |
| NSC | No Substrate Control | Blank | N/A | N/A |
| Lax | Laxative | Polyethylene glycol (PEG) 3350 | 17 | N/A |
| PH | Psyllium Husk | Single fiber | 11.6 | 6 |
| B3 | 3-Fiber Blend | Blend of resistant potato starch, resistant maltodextrin, and stabilized rice bran | 9.2 | 5.5 |
| B5 | 5-Fiber Blend | Blend of B3 ingredients + partially hydrolyzed guar gum, soluble corn fiber | 11.5 | 7.5 |
| B5 + AO | 5-Fiber + Antioxidant | B5 + Green Tea Extract | 11.7 | 7.5 |
| B5 + Bar | 5-Fiber + Bar | B5 + whole-food bar matrix | 48* | 11<sup>#</sup> |

\*: Dose reflects total mass of bar formulation added to match a single serving equivalent of the 5-Fiber Blend.
<sup>#</sup>: Fiber content of B5+Bar is composed of 7.5g of B5 with 3.5g fiber provided by whole-food ingredients (see **Supplemental Table S2**).

### Systemic Intestinal Fermentation Research (SIFR®) Technology

First, gastric and small intestinal digestion and absorption simulation was performed via the modified INFOGEST 2.0 protocol, described by Van den Abbeele, et al.^41–43^ Due to its small molecular size, the osmotic laxative (Lax) was not subject to upper gastrointestinal simulation, as it would be removed during dialysis steps designed to eliminate digested products such as amino acids (from proteins) and sugars (from starch or maltodextrin). Instead, Lax was added directly at the start of the colonic incubations to ensure full delivery to the colon, consistent with its *in vivo* mode of action. Following upper gastrointestinal simulation, test products were introduced into colonic fermentation reactors containing fecal microbiota from PD patients. Colonic simulation was performed as previously described. ^41,43^ Briefly, individual bioreactors were processed in parallel in a bioreactor management device (Cryptobiotix, Ghent, Belgium). Each bioreactor contained 5 mL of nutritional medium-fecal inoculum blend supplemented with digested test products derived from the simulated small intestinal digestion protocol, then sealed individually, before being rendered anaerobic. Blend M0017 was used for preparation of the nutritional medium (Cryptobiotix, Ghent, Belgium). After preparation, bioreactors were incubated under continuous agitation (140 rpm) at 37°C (MaxQ 6000, Thermo Scientific, Thermo Fisher Scientific, Merelbeke, Belgium). Upon gas pressure measurement in the headspace, liquid samples were collected for subsequent analysis.

### Key Fermentation Parameters

Fermentation was evaluated by production of short-chain fatty acids (SCFA), branched chain fatty acids (BCFA), pH, and gas production, as previously described.^43^ Briefly, SCFAs (acetate, propionate, butyrate, and valerate) and BCFAs (isobutyrate, isocaproate, and isovalerate) were determined via GC with flame ionization detection (Trace 1300, Thermo Fisher Scientific, Merelbeke, Belgium), upon diethyl ether extraction.^43,44^ pH was measured using an electrode (Hannah Instruments Edge HI2002, Temse, Belgium).

### Microbial Composition Analysis

The composition of the fecal microbiomes was evaluated at baseline (0h) and then after 24h incubation with each test product via quantitative shallow shotgun sequencing. Quantitative insights were obtained by correcting proportions (%) with total counts (cells/ml; flow cytometry), resulting in estimated cells/mL of different taxa. First, to determine total counts, samples were diluted in anaerobic phosphate-buffered saline (PBS), followed by cell staining with SYTO 16 at a final concentration of 1μM, and counted via a BD Novocyte Quanteon (Agilent). Data were analyzed using NovoExpress, version 1.6.2. DNA was extracted via the SPINeasy DNA Kit for Soil (MP Biomedicals, Eschwege, Germany), according to manufacturer’s instructions. Subsequently, a library was prepared using the Nextera XT DNA Library Preparation Kit (Illumina, San Diego, CA, USA) and IDT Unique Dual Indexes (total DNA input, 1 ng). A proportional amount of Illumina Nextera XT fragmentation enzyme was added to fragment genomic DNA. Libraries were constructed, purified, and quantified as previously described^43^, then sequenced on an Illumina Nextseq 2000 platform 2 × 150 base pairs. The CosmosID-HUB Microbiome Platform (CosmosID Inc., Germantown, MD, USA) was used to convert unassembled sequencing reads to relative abundances (%) of taxa.^45,46^

### Microbial Metabolite Quantification

Metabolomics analysis was performed as previously described.^47,48^ In brief, the analysis was carried out using a Vanquish UHPLC (Thermo Scientific, Germering, Germany) coupled to Orbitrap Exploris 240 MS (Thermo Scientific, Bremen, Germany) using an electrospray ionization source, applied both in negative and positive ionisation mode. The UPLC was performed using an adapted version of the protocol described previously.^49^ Peak areas were extracted using Compound Discoverer 3.3 (Thermo Scientific), along with a manual extraction based on an in-house library using Skyline 24.1 (MacCoss Lab Software).^50^ Technical variability was confirmed by running a quality control sample (pooled aliquots of all samples) every six samples.

A total of 525 compounds were detected at 24 hours and annotated across four levels of identification confidence. Annotation levels were defined as: Level 1, confirmation by retention time matched to in-house authentic standards, accurate mass (≤3 ppm deviation), and MS/MS spectra; Level 2a, retention time and accurate mass; Level 2b, accurate mass and MS/MS spectra; and Level 3, accurate mass alone. Given the higher degree of confidence in Level 1 and 2a annotations, subsequent analyses focused on Level 1/2a metabolites. A complete list of detected metabolites, including annotation levels, is provided in **Supplemental File 1.**

### Data Analysis and Statistics

Statistical analyses were conducted using GraphPad Prism (version 10.6.1; GraphPad Software, San Diego, CA, USA) and Microsoft Excel (Microsoft Corp., Redmond, WA, USA). Donor-matched comparisons across treatment conditions were evaluated using repeated-measures one-way analysis of variance (ANOVA), with donor treated as a matched factor. When sphericity could not be assumed, the Geisser-Greenhouse correction was applied. Post hoc comparisons between column means were performed using Tukey’s multiple comparisons test.

## Results

### Baseline Inflammatory and Microbial Heterogeneity Across Parkinson’s Disease Donors

Fecal calprotectin and zonulin were measured from donor stool to assess intestinal inflammation and gut barrier integrity, respectively **(Figure 1A, B).** Using established cutoffs for inflammation status, most donors fell within the non-inflamed range (11-53 μg/g), whereas one donor exhibited moderate intestinal inflammation (167 μg/g) **(Figure 1A)**. Zonulin regulates epithelial tight junctions and is a marker of intestinal barrier integrity.^51^ Fecal levels below 110 ng/g are considered within the healthy range, whereas elevated levels may indicate increased permeability.^52^ All donors fell below this cutoff, with values ranging from 25-97 ng/g **(Figure 1B).** Notably, zonulin has also been implicated in regulation of blood-brain barrier permeability, linking intestinal and neurological barrier function in neurodegenerative disease.^53–55^

**Figure 1:**
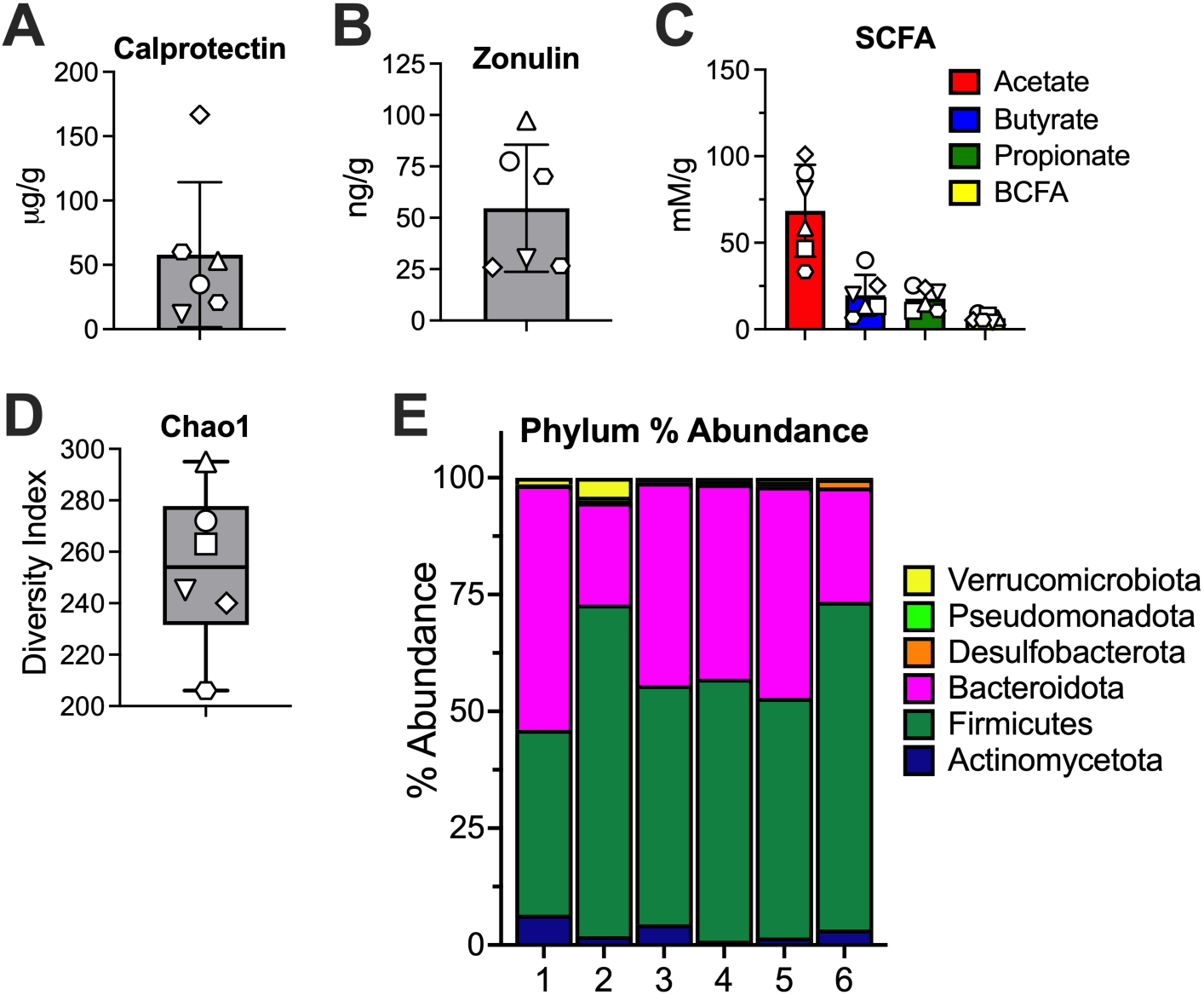
Baseline inter-individual heterogeneity in inflammatory, metabolic, and microbial features of Parkinson’s disease (PD) donor microbiomes. Baseline (0h) donor stool samples were analyzed for: **A)** fecal calprotectin concentrations (μg/g fecal sample), **B)** fecal zonulin concentrations (ng/g fecal sample), **C)** short-chain fatty acid (SCFA; acetate, butyrate, propionate) and branched-chain fatty acids (BCFA) concentrations (mM/g fecal sample), **D)** alpha diversity assessed by the Chao1 index, and **E)** relative (%) abundance of bacterial phyla. Bars represent the mean +/- standard deviation (SD) **(A, B, C)**. Box-and-whisker plots show the median (center line), interquartile range (box), and minimum and maximum values (whiskers) **(D)**. Individual donors are denoted by unique symbols: donor 1, circle; donor 2, square; donor 3, triangle; donor 4, inverted triangle; donor 5, diamond; donor 6, hexagon.

Baseline fermentation parameters were assessed by measuring pH, SCFAs, and BCFAs prior to simulated colonic incubation. Baseline pH values were similar across donors, indicating comparable physicochemical conditions prior to incubation **(Supplemental Figure S1A).** SCFA and BCFA concentrations per gram of fecal sample were measured at baseline **(Figure 1C).** No gas production was detected prior to fermentation, as expected. These low baseline metabolite levels establish a functionally substrate-limited state, allowing donor-specific fermentation responses to experimental formulations to be evaluated without confounding from pre-existing metabolic activity.

Microbial community composition was assessed by shallow shotgun metagenomic sequencing. α-diversity metrics revealed marked inter-donor variability in both species richness (Chao1) **(Figure 1D)** and evenness (Shannon and reciprocal Simpson indices)^56^ **(Supplemental Figure S1B, C)**. Total bacterial cell counts also differed substantially between donors **(Supplemental Figure S1D, E).** At the phylum level, all donors were dominated by Bacteroidota (21–52%) and Firmicutes (39–70%) **(Figure 1E)**, consistent with typical adult gut microbiomes.^57^ However, substantial variation was observed in lower abundance phyla, including Actinomycetota (0.9–6.5%), Desulfobacterota (0.1–1.8%), Pseudomonadota (0.06–0.9%), and Verrucomicrobiota (0.009–4%), taxa that include organisms involved in fermentation, mucin degradation, and inflammatory signaling. Composition at the family level further illustrates the differences between donors, with varying abundances of *Bacteroidaceae* (18-48%), *Ruminococcaceae* (5-18%), *Lachnospiraceae* (22-37%) **(Supplemental Figure S1F).** There were also stark differences in the presence and abundance of potentially pathogenic bacteria in the donors, with *Akkermansiaceae* ranging from 0.009-4% abundance, and *Enterobacteriaceae* ranging from 0.003-0.07% **(Supplemental Figure S1F).** Collectively, these profiles demonstrate that the PD donor microbiomes used in this study differ widely in both composition and functional potential, underscoring the need to evaluate formulation effects across diverse microbial ecosystems rather than relying on a single representative microbiome.

### An Osmotic Laxative Does Not Significantly Alter Microbial Fermentation or Community Structure

Because osmotic laxatives are commonly used to manage constipation in Parkinson’s disease, we first examined whether an osmotic Laxative (Lax, light gray) alters microbial fermentation relative to the No Substrate Control (NSC, black). Total SCFA production, individual SCFAs, culture pH, and gas production were used as primary readouts of microbial metabolic engagement **(Figure 2 and Supplemental Figure S2).** Laxative treatment did not increase total SCFA production **(Figure 2A)** or concentrations of acetate, butyrate, or propionate relative to NSC **(Figure 2B-D).** Culture pH remained comparable between Lax and NSC conditions **(Figure 2E),** consistent with minimal generation of fermentation-derived organic acids. Gas production was not significantly changed relative to NSC **(Figure 2F and Supplemental Figure S2B).** These findings confirm that polyethylene glycol-based osmotic laxatives do not provide fermentable substrate capable of supporting microbial metabolic activity within PD donor microbiomes.

**Figure 2:**
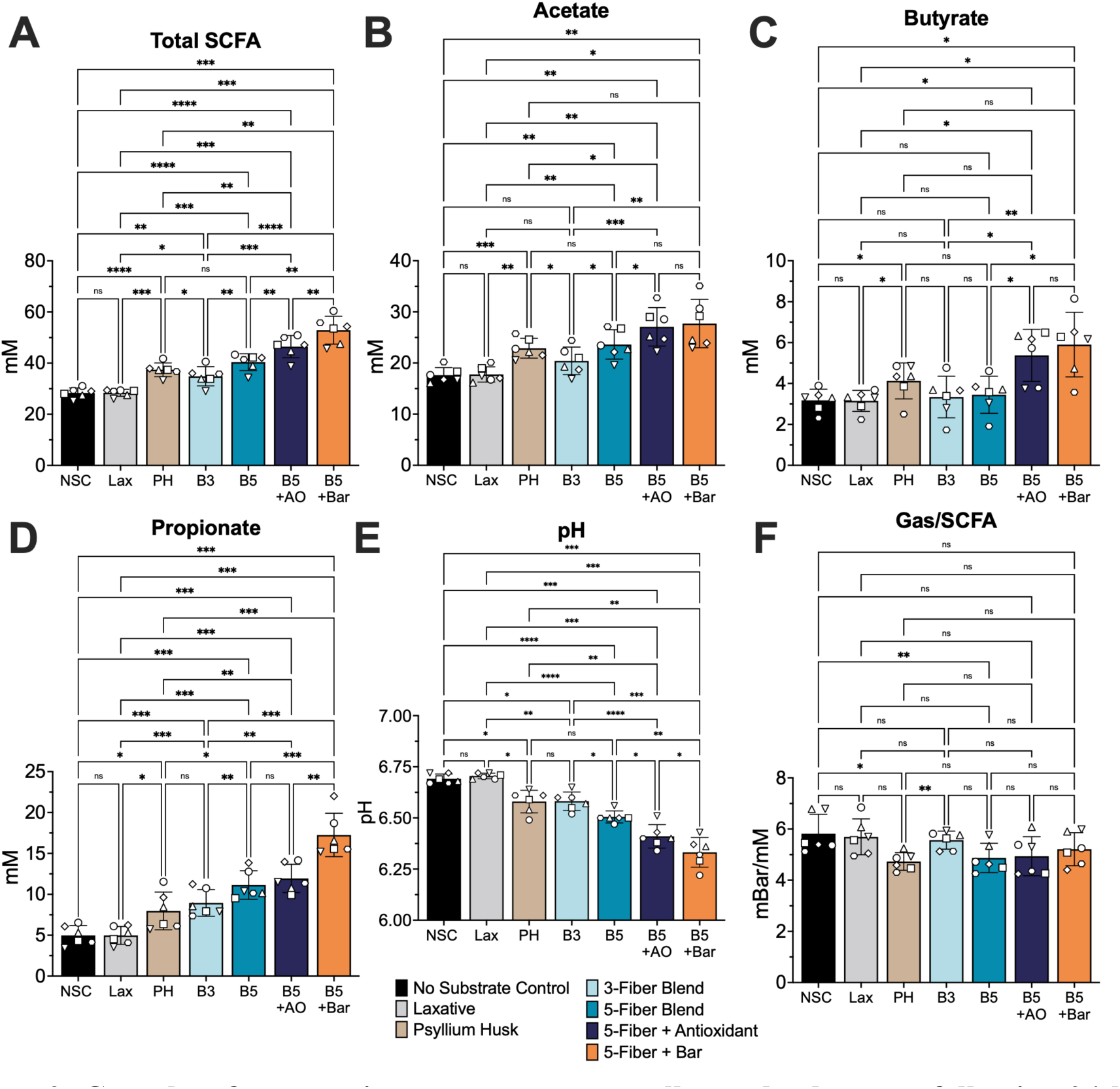
Complete fermentation outcomes across all tested substrates following 24-hour anaerobic ex vivo fermentation. Donor stool samples were fermented anaerobically for 24 h with each test product. Fermentation outcomes included: **A)** total SCFA (mM; acetate + butyrate + propionate + valerate), **B)** acetate (mM), **C)** butyrate (mM), **D)** propionate (mM), **E)** pH, **F)** gas per SCFA ratio (mBar/mM). Test products are color-coded as follows: No Substrate Control (NSC), black; Laxative (Lax), light gray; Psyllium Husk (PH), light brown; 3-Fiber Blend (B3), light teal; 5-Fiber Blend (B5), dark teal; 5-Fiber Blend + Antioxidant (B5+AO), dark blue; 5-Fiber Blend + Bar excipients (B5+Bar), orange. Bars represent the mean +/- standard deviation (SD) **(A-F).** Individual donors are represented by symbols as in Figure 1. Statistical significance was evaluated using repeated-measures one-way ANOVA with Geisser-Greenhouse correction followed by Tukey’s multiple comparisons tests. ns, non-significant; *, p < 0.05; **, p < 0.01; ***, p < 0.005; ****, p < 0.0001.

### Increasing Substrate Complexity Enhances SCFA Production Across PD-Derived Microbiomes

We next examined how fiber formulations can reshape microbial metabolism. We compared a single fiber (Psyllium Husk, PH, tan) with increasingly complex fiber blends (3-Fiber Blend, B3, light teal; 5-Fiber Blend, B5, teal) to assess how substrate architecture shapes microbial fermentation and SCFA profiles **(Figure 2).** As expected, fermentable substrates increased total SCFA production relative to the NSC with the highest overall SCFA levels observed for B5 **(Figure 2A).** PH increased acetate production relative to NSC and B3, while incorporation of additional fiber substrates in B5 increased acetate levels to approximately the same as Psyllium Husk **(Figure 2B).** Butyrate production increased slightly with fiber treatment but showed more variable responses between donors **(Figure 2C).** Propionate production increased with increasing number of fiber sources **(Figure 2D).** BCFA production was low across all conditions, with B5 showing the lowest levels, suggesting a relative shift toward saccharolytic rather than proteolytic fermentation **(Supplemental Figure S2A).**

In addition to SCFA production, pH and gas output provided complementary readouts of microbial fermentation. Fiber substrates progressively lowered culture pH relative to the NSC, with B5 producing the greatest acidification compared to PH and B3 (mean 6.51), consistent with enhanced fermentative activity **(Figure 2E).** Although gas production increased with substrate complexity **(Supplemental Figure S2B),** the ratio of gas/SCFA produced for B5 was comparable to that of PH and lower than that of B3 **(Figure 2F).**

Given the heterogeneity of PD donor microbiomes, formulations containing multiple complementary fiber substrates may be better suited to support consistent microbial metabolic engagement than single-fiber interventions.^58^

### Formulation Context modulates SCFA production from a core fiber blend

Having identified B5 as the formulation resulting in the greatest overall SCFA levels, we next evaluated how modifications to this core substrate can further alter fermentation parameters. We tested the 5-Fiber Blend supplemented with an antioxidant (5-Fiber + Antioxidant, B5+AO, dark blue) or delivered within a whole-food bar matrix (5-Fiber + Bar, B5+Bar, orange) to assess how formulation context influences SCFA production **(Figure 2).** As expected for a formulation providing fermentable substrate within a whole-food matrix, the B5+Bar condition supported the greatest total SCFA production among fiber-containing treatments **(Figure 2A).** Incorporation of a natural antioxidant into the fiber mixture was associated with increased butyrate production relative to the base B5 treatment **(Figure 2C).** These formulation modifications also increased propionate production **(Figure 2D)**. pH inversely correlated with total SCFA production, with the bar matrix resulting in the most acidic environment **(Figure 2E).** The bar also led to the most gas produced **(Supplemental Figure S2B);** however, the ratio of gas to SCFA produced remained comparable to the other formulations, indicating efficient conversion of substrate into microbial metabolites **(Figure 2F).**

Across PD donor microbiomes, incorporation of antioxidant supplementation or delivery within a whole-food matrix was associated with increased SCFA production relative to the base 5-Fiber Blend. These findings indicate that non-fiber formulation components can further augment microbial fermentative activity, providing additional strategies to optimize prebiotic interventions.

### Fiber formulations remodel microbial community structure in a donor-dependent manner

To determine whether the observed fermentation-associated changes were accompanied by changes in microbial community structure and composition, we performed microbiome profiling using shallow shotgun metagenomic sequencing before and after 24 h anaerobic *ex vivo* fermentation with each formulation **(Figure 3).** Principal coordinates analysis (PCoA) based on Bray-Curtis dissimilarity of bacterial family-level relative abundances revealed distinct clustering by substrate type **(Figure 3A).** The NSC and Lax conditions clustered closely together, consistent with minimal fermentative or community-level effects, whereas fiber-containing formulations diverged from these controls. B5 and B5+AO clustered together but were distinctly separate from B5+Bar, indicating that the bar matrix ingredients drive microbiome remodeling distinctly from the base fiber blend.

**Figure 3.**
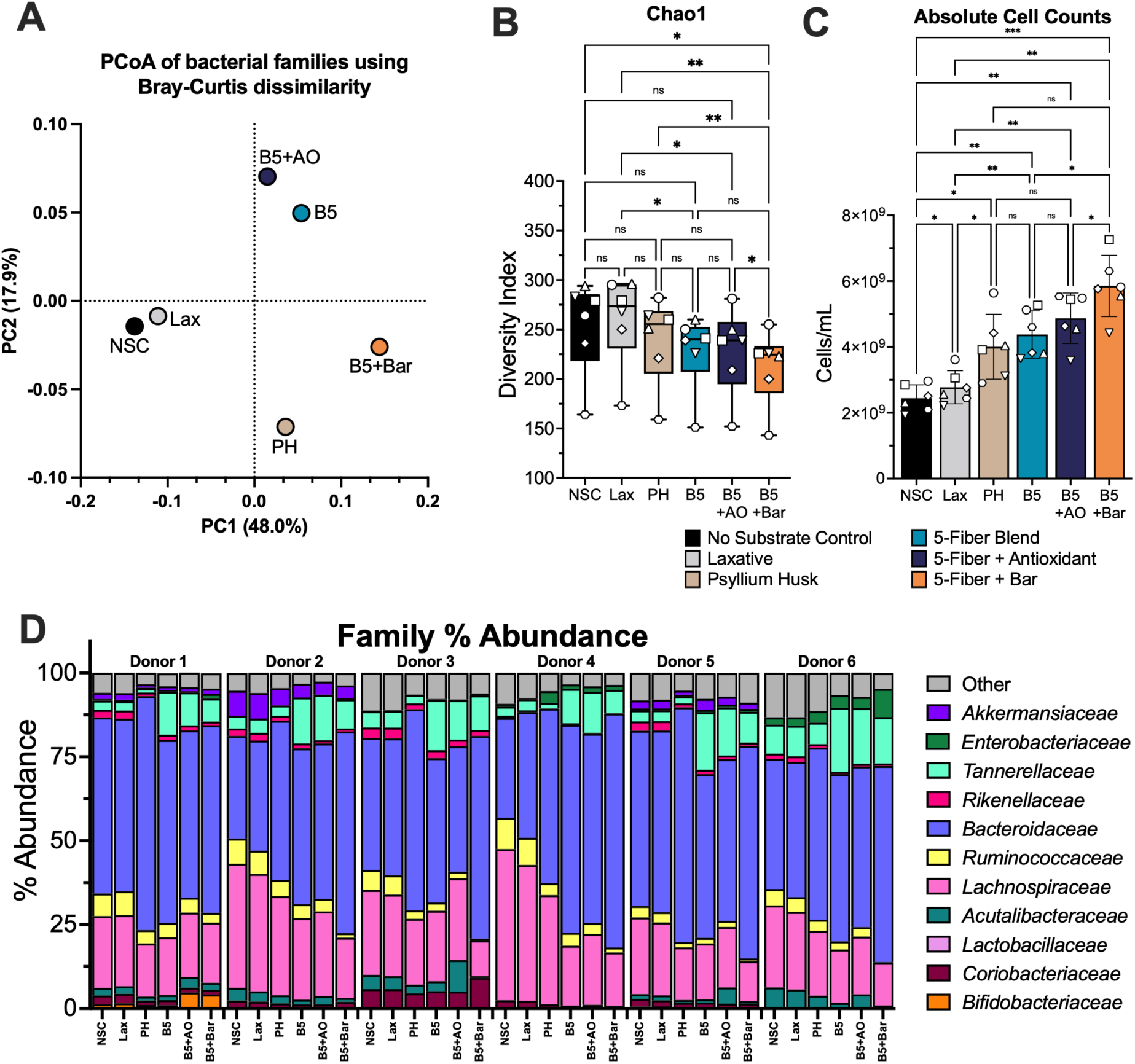
Fiber blend complexity and formulation modulate microbiome diversity, community structure, and taxonomic composition following ex vivo fermentation. Following 24 h of anaerobic ex vivo fermentation, microbiome outcomes were assessed: **A)** principal coordinates analysis (PCoA) based on Bray-Curtis dissimilarity of bacterial family-level relative abundances, **B)** alpha diversity assessed by the Chao1 index, **C)** averaged absolute bacterial cell counts, and **D)** relative abundance (%) of top bacterial families. Bars represent the mean ± SD **(C).** Box-and-whisker plots show the median (center line), interquartile range (box), and minimum and maximum values (whiskers) **(B).** Individual donors are represented by symbols as defined in Figure 1. Statistical significance was assessed using repeated-measures one-way ANOVA with Geisser–Greenhouse correction followed by Tukey’s multiple comparisons test. ns, non-significant; *, p < 0.05; **, p < 0.01; ***, p < 0.005; ****, p < 0.0001.

Alpha diversity metrics exhibited modest but consistent downward trends with fiber-containing treatments, as assessed by Chao1 **(Figure 3B)**, Shannon, and reciprocal Simpson indices **(Supplemental Figure S3A, B).** This pattern is consistent with selective enrichment of fiber-responsive taxa rather than broad community expansion and aligns with prior studies reporting reduced alpha diversity following dietary fiber interventions.^59,60^ In contrast to diversity metrics, total bacterial biomass increased significantly with fiber-containing substrates across all donors, as reflected by increased absolute bacterial cell counts **(Figure 3C)**, indicating enhanced microbial growth despite reduced within-sample diversity.

Donor-specific differences in microbiome composition persisted following substrate exposure. At higher taxonomic levels, relative abundance profiles varied substantially between donors across all conditions **(Figure 3D and Supplemental Figure S3C)**, highlighting pronounced inter-individual heterogeneity that was maintained even in the presence of fermentable substrates. Given this, taxon-level responses were evaluated across donor-specific ranges rather than relying solely on group means **(Figure 4).** Following incubation under NSC conditions, *Bacteroides* abundance spanned a wide range across donors (4-43%) **(Figure 4A).** Laxative treatment resulted in a nearly identical distribution, consistent with its lack of fermentable activity. In contrast, fiber-containing formulations expanded the upper range of *Bacteroides* abundance across donors. Notably, PH treatment yielded the broadest expansion, with *Bacteroides* relative abundance reaching up to ∼54% in high-responder donors **(Figure 4A).**

**Figure 4.**
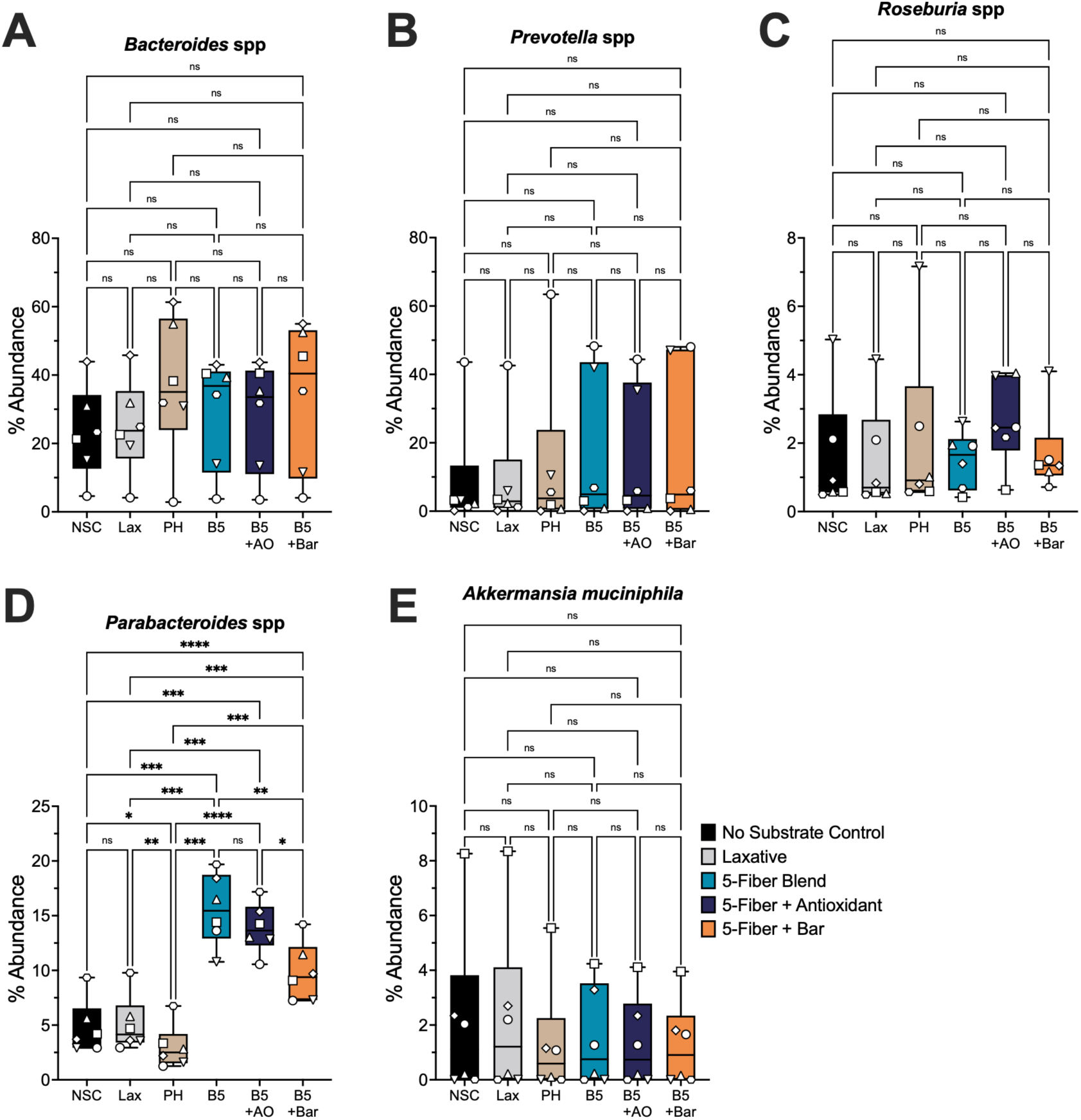
Fiber blend complexity and formulation alter the relative abundance of select bacterial genera and species following ex vivo fermentation. Relative abundance (%) of A) *Bacteroides* species, B) *Prevotella* species, C) *Roseburia* species, D) *Parabacteroides* species, and E) *Akkermansia muciniphila* were quantified following 24 h of anaerobic ex vivo fermentation. Box-and-whisker plots show the median (center line), interquartile range (box), and minimum and maximum values (whiskers) **(A-E).** Individual donors are represented by symbols as defined in Figure 1. Statistical significance was assessed using repeated-measures one-way ANOVA with Geisser–Greenhouse correction followed by Tukey’s multiple comparisons test. ns, non-significant; *, p < 0.05; **, p < 0.01; ***, p < 0.005; ****, p < 0.0001.

Other genera exhibited similarly donor-dependent but substrate-associated trends. *Prevotella* species **(Figure 4B)** and *Roseburia* species **(Figure 4C),** taxa frequently reported as depleted in PD,^21,61^ showed directional increases with fiber-containing formulations in subsets of donors. *Parabacteroides* species exhibited the most consistent treatment-associated increase among the taxa examined, reaching statistical significance in select comparisons **(Figure 4D).** *Akkermansia muciniphila* relative abundance displayed variable levels across donors but showed a general decrease in abundance with fiber products compared to NSC or Lax **(Figure 4E).**

Collectively, these data demonstrate that fiber blend complexity and formulation modulate microbial community structure primarily through increased biomass and selective expansion of responsive taxa, with taxon-level outcomes shaped by substantial donor-specific heterogeneity rather than uniform shifts across all microbiomes.

### Formulation-dependent shifts in microbial metabolic pathways beyond SCFAs

Short-chain fatty acids are not the only metabolites produced by the gut microbiome.^62,63^ To further characterize the functional consequences of fiber blend complexity and formulation, we evaluated the broader metabolite pool following 24 h anaerobic *ex vivo* fermentation **(Figure 5),** focusing on metabolites implicated in gut-brain axis signaling and those previously reported to differ in Parkinson’s disease. Metabolite abundances are presented as a heat map of log2 fold change relative to NSC and averaged across six donors; values exceeding the visualization limits of the heat map (± 4 log2FC) are indicated with a hash symbol (#) with full fold-change distributions provided in **Supplemental Figure S4.**

**Figure 5:**
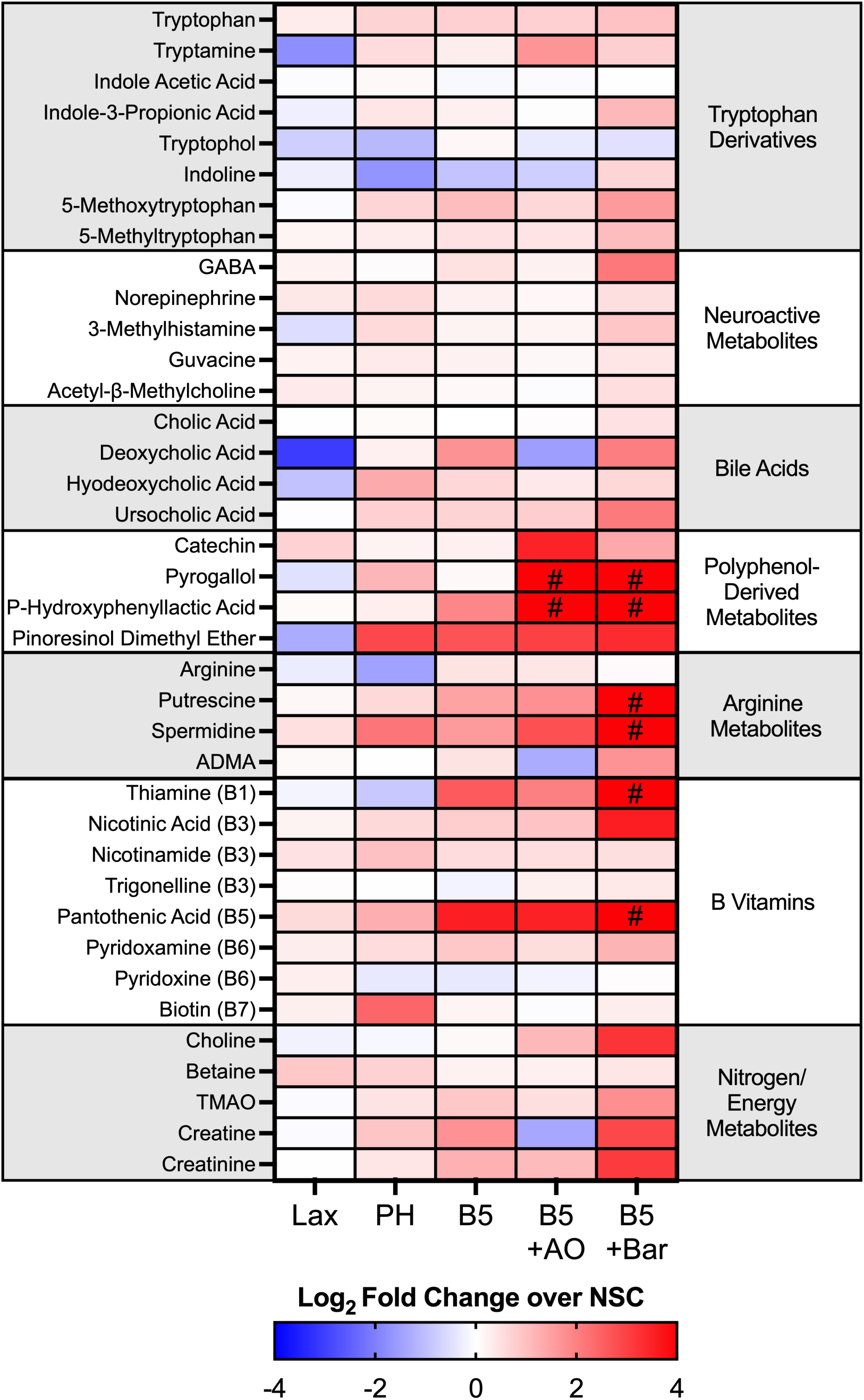
Fiber blend complexity and formulation modulate gut-brain axis-relevant metabolic outputs following ex vivo fermentation. Untargeted metabolomic profiling was performed on fermentation supernatants following 24 h anaerobic ex vivo fermentation. Heat maps display the log_2_ fold change (log_2_FC) in metabolite abundance for each test product relative to the no-substrate control (NSC). Metabolites are grouped by functional relevance in the following order: tryptophan derivatives, neuroactive metabolites, primary and secondary bile acids, polyphenol-derived metabolites, arginine metabolites, B vitamins, and nitrogen/energy metabolism-related compounds. Color intensity represents the magnitude and direction of change relative to NSC, with red indicating increased abundance and blue indicating decreased abundance. Hash symbols (#) denotes metabolites exhibiting high-magnitude increases (> +4 log_2_FC) relative to NSC.

Across conditions, fiber-containing substrates elicited coordinated, pathway-level shifts in microbial metabolites relative to the NSC, with the magnitude and direction of change varying by fiber blend complexity and formulation. Within tryptophan metabolism, fiber exposure was associated with directional changes in indole-derived metabolites, indicating altered microbial processing of tryptophan relative to NSC **(Figure 5).** Neuroactive metabolites similarly exhibited formulation-dependent shifts with fiber-containing conditions, consistent with altered microbial contributions to neuroactive metabolite pools.

Bile acid-associated metabolites displayed selective, formulation-dependent responses **(Figure 5).** The primary bile acid cholic acid showed minimal change across conditions, whereas secondary bile acids were more responsive to fiber exposure and formulation. Several secondary bile acids increased with the B5 and B5+Bar formulations but were reduced under Lax and B5+AO conditions, consistent with differential modulation of microbial bile acid transformation pathways.

Polyphenol-derived metabolites exhibited some of the most robust pathway-level responses, with B5+AO and B5+Bar generally amplifying metabolite abundance relative to B5 alone, with several compounds showing high-magnitude increases (> +4 log_2_FC) relative to NSC **(Figure 5 and Supplemental Figure S4A, B).** These patterns are consistent with enhanced microbial processing of polyphenol substrates when layered onto a multi-fiber background. Metabolites related to arginine metabolism, including polyamines, were increased with fiber-containing conditions and were most pronounced in formulations containing bar excipients or antioxidant supplementation (**Figure 5 and Supplemental Figure S4C, D)**. B vitamin-associated metabolites also showed enrichment across fiber-containing treatments, particularly with B5+Bar, with select vitamins increased across all 5-Fiber Blend-containing products **(Figure 5 and Supplemental Figure S4E, F).**

Finally, metabolites related to nitrogen and energy metabolism exhibited formulation-dependent modulation, particularly in formulations containing increased fiber complexity and peaking with additional formulation components. Levels of choline, a precursor for acetylcholine, increased ∼2 log_2_FC with B5+AO and almost 3 log_2_FC with B5+Bar **(Figure 5).** Creatine and creatinine were similarly elevated in the bar condition, whereas creatine levels were reduced in the B5+AO formulation. While individual metabolite responses were donor-dependent, consistent pathway-level patterns across multiple metabolite classes support the conclusion that fiber blend complexity and formulation features modulate microbial metabolic capacity beyond SCFA production.

## Discussion

The gastrointestinal tract and its resident microbiota are increasingly recognized as key regulators of host physiology, including neurological health. The gut microbiome influences immune function, endocrine and metabolic signaling, and neural communication, contributing to dynamic interactions between the enteric and central nervous systems in both health and disease. Accordingly, the role of the gut microbiota in neurodegenerative disorders, such as Parkinson’s disease (PD) or Alzheimer’s disease (AD)^64–66^, has become an active and rapidly evolving area of research. Multiple preclinical and clinical studies have identified altered microbial community structures and metabolite profiles in PD, giving rise to “brain-first” and “body-first” models of disease onset that incorporate microbiome-related mechanisms, including hypotheses in which pathological processes may originate in the gastrointestinal tract and propagate to the central nervous system.^17,61,67–69^ This expanding body of work has fueled growing interest in microbiome-targeted interventions as tools to better understand and potentially influence PD pathophysiology.

Among these approaches, dietary fibers represent an attractive strategy due to their capacity to modulate microbial metabolism through provision of fermentable substrates. However, accumulating evidence suggests that fiber source, physicochemical structure, and fermentative accessibility are critical determinants of microbial response.^70,71^ Not all fibers elicit equivalent ecological or metabolic effects, and inter-individual differences in microbial enzymatic capacity further shape functional outcomes.^58,59^ In this study, we employed the *ex vivo* SIFR^®^ technology platform, which has been shown to generate clinically relevant insights into both microbiome modulation and host-relevant functional responses.^41,43^ We identified that both the number of fibers within a blend and the inclusion of non-fiber additives substantially influenced microbial composition and metabolic outputs. By systematically varying substrate complexity and formulation context, we demonstrate that fiber architecture shapes microbial fermentation capacity across heterogeneous PD-derived microbiomes **(Figure 2).**

An important design feature of the present study is the inclusion of a clinically relevant osmotic laxative comparator. Few microbiome-directed dietary intervention studies directly contrast fermentable substrates with standard, non-fermentable, therapies used to manage constipation. Incorporating polyethylene glycol enables mechanistic differentiation between luminal water–mediated stool softening and fermentation-driven modulation of the intestinal ecosystem. As expected, based on its mechanism of action^72^, polyethylene glycol did not significantly alter SCFA production or microbial community structure in the present study **(Figure 2 and Supplemental Figure S3).** In contrast, fermentable fiber formulations shifted microbial metabolism toward saccharolytic fermentation and increased SCFA output. This comparison underscores an important distinction between symptom-focused interventions that modify stool consistency and substrate-based strategies that engage microbial metabolic pathways. In PD populations, where reduced SCFA-producing taxa and altered microbial function have been reported, interventions that directly stimulate fermentative metabolism may represent a complementary approach to managing gastrointestinal dysfunction.

Importantly, the results of this study further underscore that “fiber” is not a uniform intervention. Fermentability depends on physicochemical structure, solubility, branching patterns, and accessibility to microbial carbohydrate-active enzymes.^70^ Individual taxa differ in enzymatic repertoires, leading to substrate-specific utilization and cross-feeding dynamics within microbial consortia, which are governed by cooperative and competitive interactions that structure polymicrobial communities.^73,74^ Increasing substrate complexity through multi-fiber blends likely broadens ecological engagement across microbial guilds, supporting more consistent SCFA production across heterogeneous PD microbiomes. Consistent with this ecological framework, microbial responses to dietary substrates are shaped not only by substrate availability but also by competitive interactions within the gut ecosystem, where organisms may utilize alternative or non-preferred substrates to maintain fitness under conditions of resource limitation.^75^ The enhanced metabolic responses observed with the 5-Fiber Blend relative to single-fiber intervention align with emerging evidence that systematically designed fiber mixtures can outperform isolated substrates in promoting coordinated microbial output. Thus, formulation architecture, encompassing complementary substrates rather than fiber quantity alone, appears central to optimizing microbiome engagement.

Because gastrointestinal discomfort is frequently reported with high-fiber supplementation, the balance between fermentative efficiency and gas production warrants consideration. Although gas production increased with substrate complexity, the ratio of gas to SCFA production remained comparable or improved in the more complex formulations, suggesting efficient substrate conversion rather than disproportionate gaseous byproducts. This distinction may be particularly relevant in PD, where altered motility and visceral sensitivity can exacerbate discomfort. Rational formulation design may therefore enhance beneficial fermentative outputs while mitigating excessive gas-related symptoms often associated with rapidly fermentable or poorly matched fibers.

Across multiple microbiome studies comparing Parkinson’s disease patients with healthy controls, a recurring pattern of altered bacterial taxa has been described, including increased abundance of *Bifidobacterium*, *Lactobacillus*, and *Akkermansia*, alongside reduced abundance of *Prevotella*, *Blautia*, and *Faecalibacterium*.^17,61,68,76^ Accordingly, a key goal of microbiome-modulating interventions in PD is to promote recovery of depleted taxa without further enriching groups that are already overrepresented. At the ecological level, fiber-containing formulations increased total microbial biomass while modestly reducing alpha diversity metrics, consistent with selective expansion of fiber-responsive taxa rather than indiscriminate community proliferation. Donor-specific variability persisted, yet directional trends were evident across formulations. Notably, *Akkermansia muciniphila*, a mucin-degrading bacterium frequently reported to be elevated in PD cohorts,^17,61,68,76^ demonstrated a general reduction in relative abundance following fiber-containing interventions. While *Akkermansia* is often considered beneficial in metabolic disease contexts, its expansion in PD and its capacity to utilize host mucus underscore the importance of disease-specific interpretation.^77^ Provision of fermentable carbohydrates may reduce reliance on host-derived mucins as carbon sources, thereby reshaping ecological niches within the intestinal environment and may have broad impacts on intestinal integrity.^78^ These findings highlight that microbial targeting strategies must account for disease context and ecological balance rather than assuming uniform benefit of individual taxa across conditions. Although intestinal inflammation has been associated with altered microbial metabolism in PD^13,14,21,39^, the present study was not powered to stratify fermentation responses by baseline inflammatory status and formulation-dependent fermentation responses were evaluated across donors irrespective of fecal calprotectin levels.

Beyond SCFAs, complex formulations elicited coordinated shifts in multiple metabolite classes implicated in host-microbe interactions. Antioxidant supplementation and whole-food matrix components amplified polyphenol-derived metabolite production, including compounds such as pyrogallol. Catechol-containing metabolites, including pyrogallol, have been reported to interact with α-synuclein and modulate fibrillization in simplified biochemical systems;^79^ however, the relevance of these observations to microbiome-derived metabolites *in vivo* remains to be established and was not assessed in the present study. Microbial conversion of polyphenols generates a range of bioactive compounds capable of influencing immune function, epithelial integrity, and redox balance.^80,81^ In parallel, additional pathway-level shifts were observed across metabolite classes including polyamines, bile acid derivatives, and B vitamin-associated metabolites. Polyamines are linked to cellular homeostasis and host-microbe interactions,^82,83^ while bile acids have been shown to participate in host signaling pathways, including immune and neuroinflammatory regulation, and can influence gut-brain axis communication.^84^ Collectively, these findings illustrate how formulation design can expand the spectrum of microbiome-derived metabolites beyond primary carbohydrate fermentation, although the functional implications of these shifts remain to be determined.

While this study provides consistent insights across multiple microbiome-derived readouts, certain considerations should be noted, including the modest number of microbiomes evaluated and the inherent variability across individuals with PD. Although sample size may appear limited (n = 6), it is important to consider the context of preclinical gut microbiome research, where studies have historically relied on single-donor models, limiting the ability to capture interpersonal variability. In contrast, the *ex vivo* SIFR^®^ technology platform enables parallel testing across multiple independent donor microbiomes under highly standardized conditions, with six donors representing a validated configuration to obtain clinically validated insights into microbiome modulation.^43^ Although larger cohorts can further strengthen statistical power, the repeated-measures design, donor-matched comparisons and convergence across fermentation chemistry, microbial biomass, community structure and metabolomics support the robustness of the findings of the present study. Future clinical studies integrating dietary intake, symptom profiling, medication use, and longitudinal microbiome analysis can further validate and extend clinically relevant insights towards *in vivo* functional outcomes.

Taken together, these findings support the concept that rational formulation design can differentially engage microbial metabolic pathways within Parkinson’s disease–associated microbiomes. By integrating complementary fermentable substrates with additional bioactive components, it may be possible to expand microbial metabolic output beyond SCFA production and influence metabolite networks implicated in both enteric physiology and neural signaling. Although conducted *ex vivo*, the consistent functional responses observed across heterogeneous donor microbiomes suggest that substrate architecture represents a modifiable lever for consistently shaping microbiome-derived mediators relevant to neurodegenerative disease. Future clinical investigations integrating gastrointestinal symptoms, microbial ecology, metabolite profiling, and neurological endpoints will be essential to determine whether targeted dietary strategies can meaningfully influence enteric–central nervous system signaling in Parkinson’s disease.

## Author Contributions

CRediT: **OAT**: Conceptualization, Formal analysis, Visualization, Writing – original draft, Writing – review & editing, Project administration. **LDV**: Methodology, Investigation, Data curation, Formal analysis, Writing – review & editing. **IAJvH**: Methodology, Investigation, Data curation, Formal analysis, Writing – review & editing. **BS**: Conceptualization, Supervision. **PVdA**: Methodology, Investigation, Data curation, Formal analysis, Writing – review & editing.

## Data Availability Statement

The datasets generated and analyzed during the current study are available from the corresponding author upon reasonable request. Processed data, including relative abundance tables and metabolite measurements supporting the findings of this study, are available and can be shared without restriction. Raw sequencing data have not been publicly deposited at this time because they are part of a larger ongoing study that includes additional proprietary formulations not reported here and are subject to pending intellectual property considerations. These constraints limit immediate public release; however, deposition in a public repository will be considered following completion of the broader study.

---

**Supplemental Figure S1:**
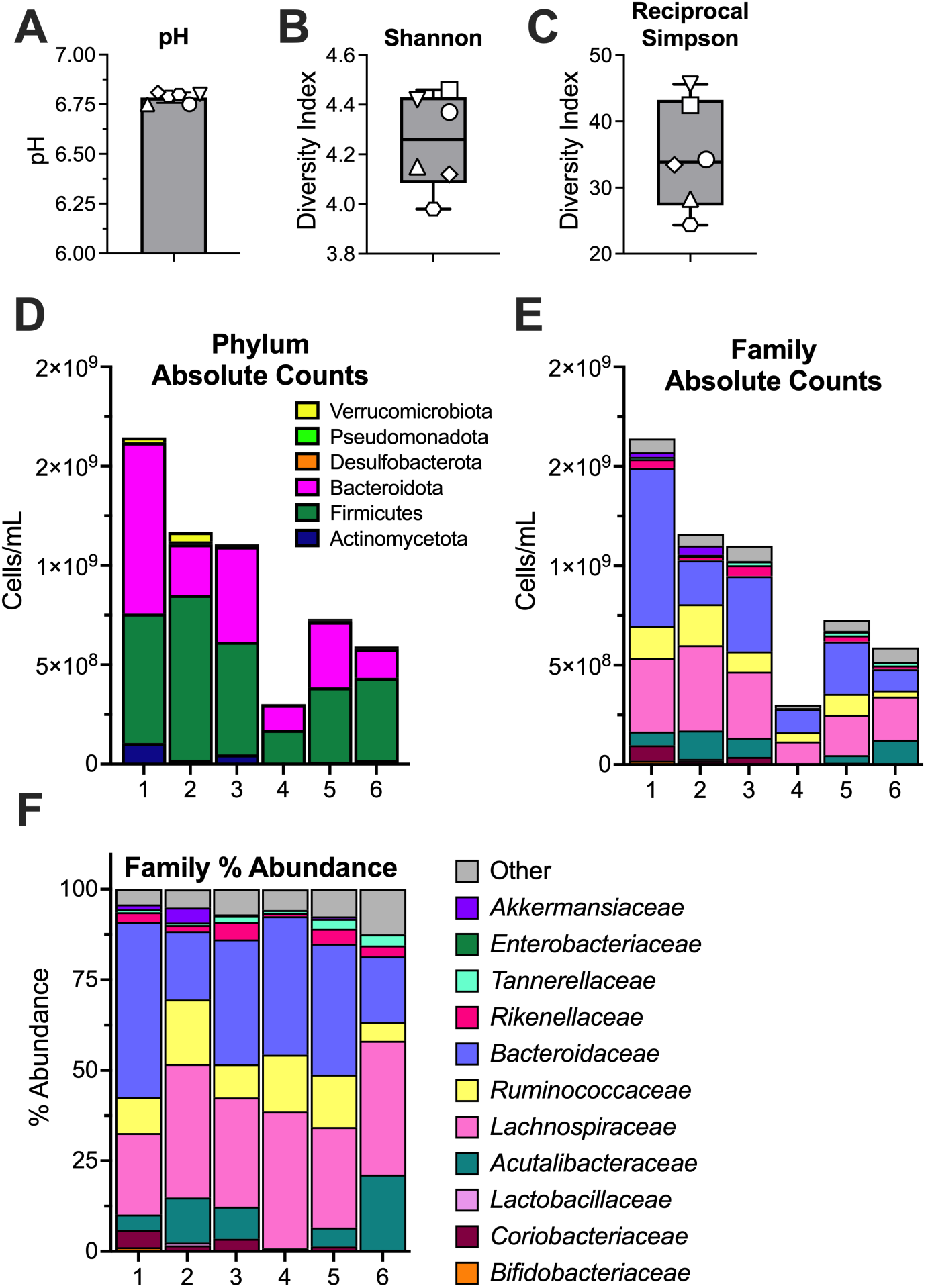
Expanded baseline metabolic and microbiome characteristics of Parkinson’s disease donor stool samples. Baseline (0 h) donor stool samples were assessed for: **A)** pH, **B)** alpha diversity metrics assessed by Shannon diversity index, **C)** alpha diversity metrics assessed by Reciprocal Simpson diversity index, **D)** absolute bacterial cell counts at the phylum level, **E)** absolute bacterial cell counts at the family level, **F)** relative (%) abundance of bacterial families. Bars represent the mean +/- standard deviation (SD) **(A).** Box-and-whisker plots show the median (center line), interquartile range (box), and minimum and maximum values (whiskers) **(B,C).** Individual donors are denoted by symbols as described in Figure 1.

**Supplemental Figure S2:**
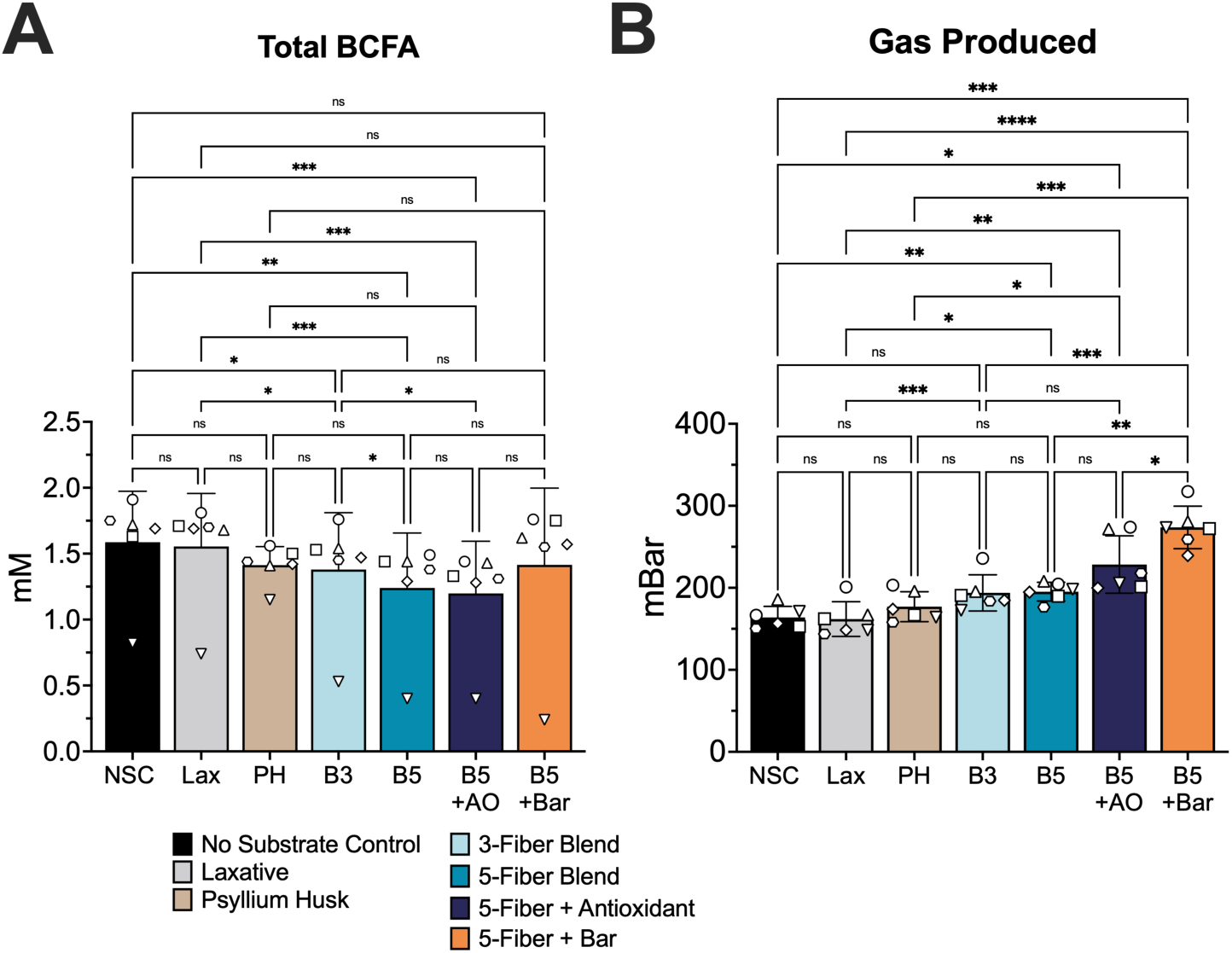
Branched-chain fatty acid production and gas output across fiber blend complexity and formulation modifications. **A)** total branched-chain fatty acids (BCFA; mM) and **B)** total gas production (mBar) Bars represent the mean ± SD **(A,B)**. Individual donors are represented by symbols as defined in Figure 1. Statistical significance was assessed using repeated-measures one-way ANOVA with Geisser–Greenhouse correction followed by Tukey’s multiple comparisons test. ns, non-significant; *, p < 0.05; **, p < 0.01; ***, p < 0.005; ****, p < 0.0001.

**Supplemental Figure S3.**
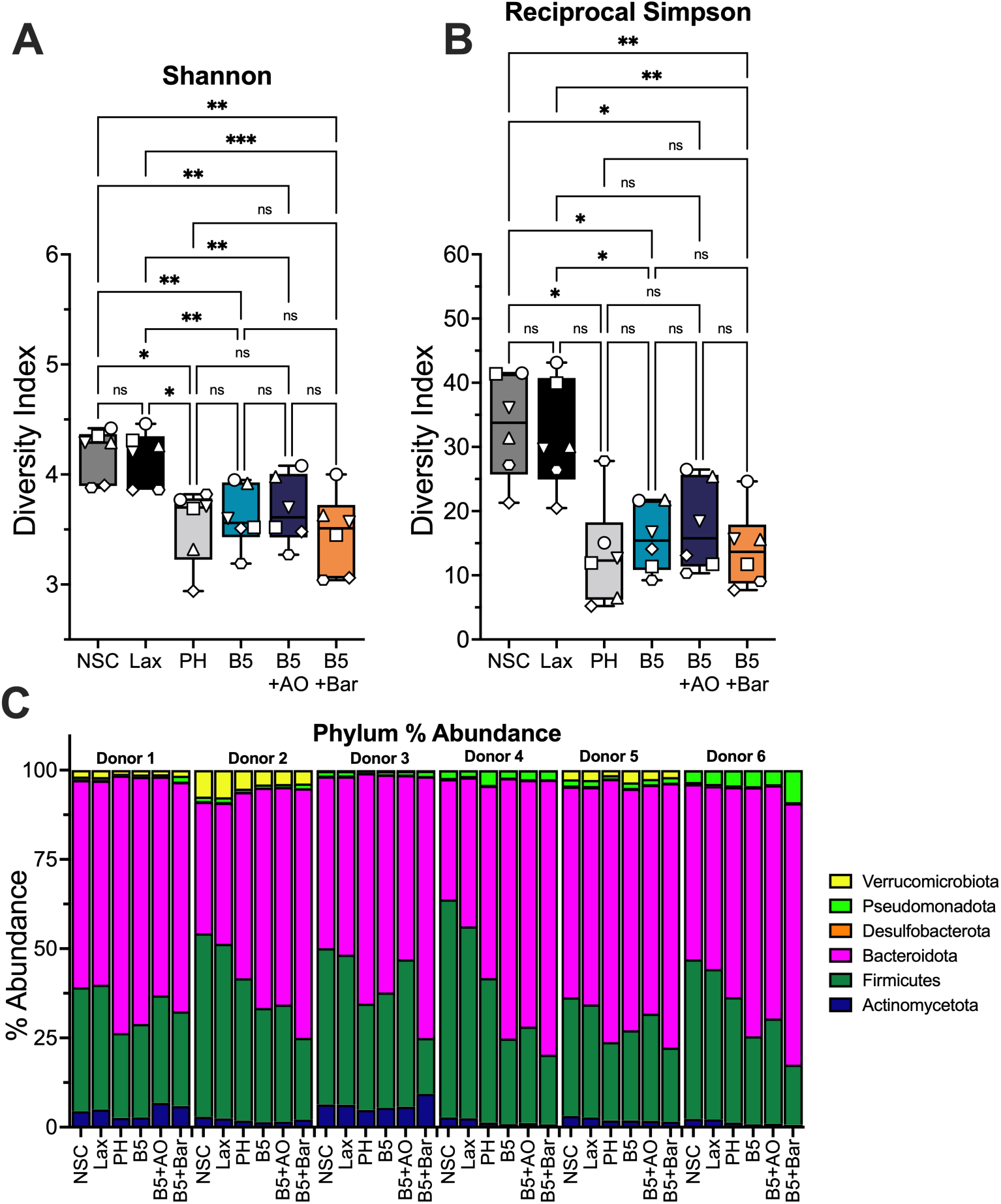
Donor-resolved microbiome diversity and composition across fermentation conditions. Following 24 h anaerobic ex vivo fermentation microbiome outcomes were assessed: **A)** alpha diversity assessed by the Shannon index, **B)** alpha diversity assessed by the reciprocal Simpson index, **C)** relative (%) abundance of bacterial phyla. Box-and-whisker plots show the median (center line), interquartile range (box), and minimum and maximum values (whiskers) (**A,B).** Individual donors are represented by symbols as defined in Figure 1. Statistical significance was assessed using repeated-measures one-way ANOVA with Geisser–Greenhouse correction followed by Tukey’s multiple comparisons test. ns, non-significant; *, p < 0.05; **, p < 0.01; ***, p < 0.005; ****, p < 0.0001.

**Supplemental Figure S4:**
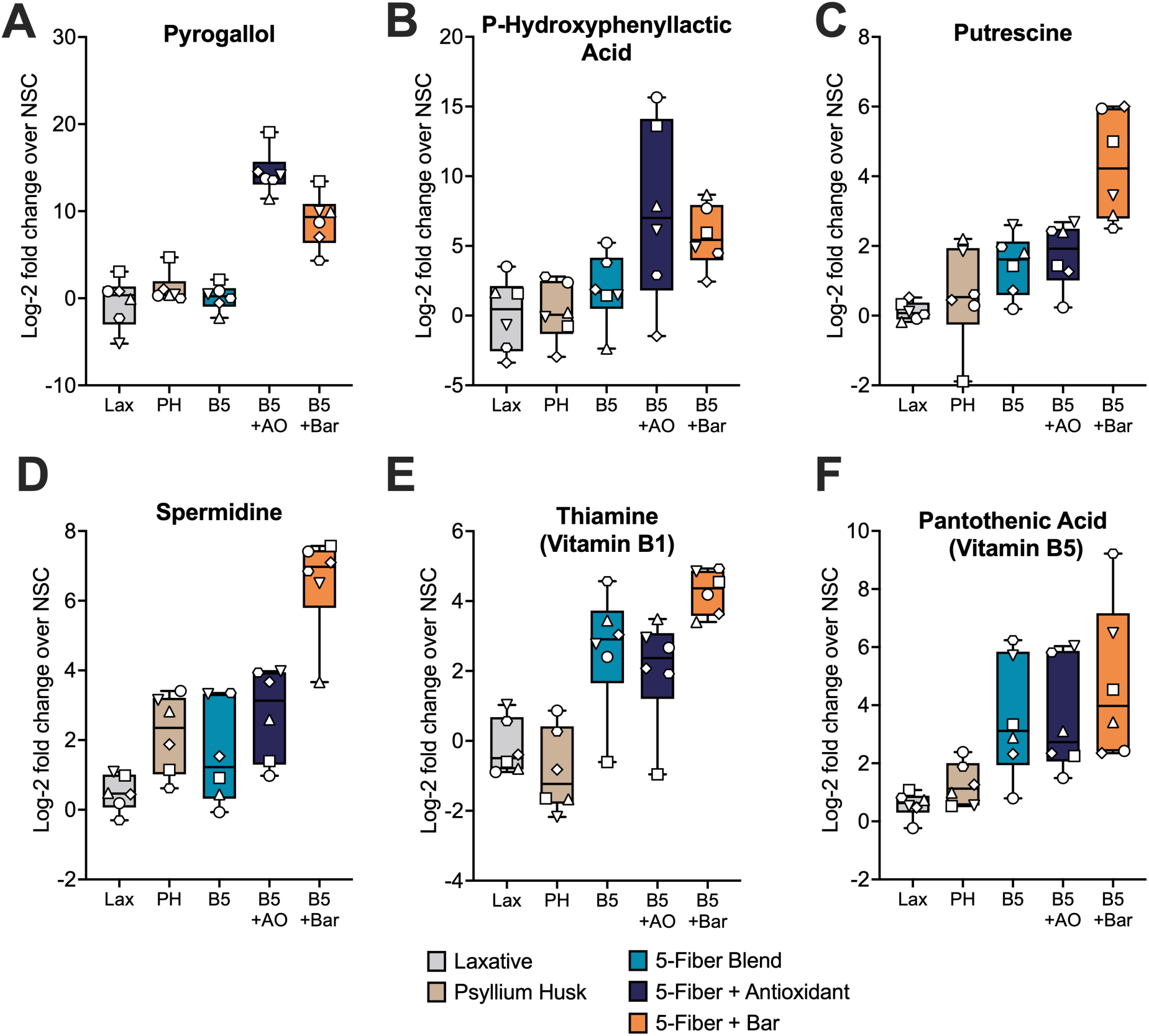
High-magnitude microbial metabolite changes following ex vivo fermentation with fiber formulations. Microbial metabolites exhibiting large-magnitude changes in abundance (log₂ fold change values exceeding ±4 relative to the No Substrate Control (NSC)) following 24 h anaerobic ex vivo fermentation: **A)** pyrogallol, **B)** p-hydroxyphenyllactic acid, **C)** putrescine, **D)** spermidine, **E)** thiamine (Vitamin B1), **F)** pantothenic acid (Vitamin B5) Box-and-whisker plots show the median (center line), interquartile range (box), and minimum and maximum values (whiskers). Individual donors are represented by symbols as defined in Figure 1.

**Supplemental Table S1:** PD Donor Characteristics. Sex, age, and medication regimen of the six microbiome donors included in the ex vivo fermentation experiments.

| **Donor** | **Sex** | **Age** | **PD Medication(s)** |
| --- | --- | --- | --- |
| 1 | F | 65 | Prolopa, Azilect |
| 2 | M | 65 | Prolopa |
| 3 | F | 55 | Prolopa, Azilect, Pramipexol |
| 4 | M | 52 | Prolopa, Pramipexol, Ecitalopram |
| 5 | F | 60 | Prolopa |
| 6 | F | 64 | Prolopa, Pramipexol, Xadago |

**Supplemental Table S2:** Ingredient-level composition and classification of test formulations used in *ex vivo* fermentation experiments.

| **Name** | **Ingredients** | **Fiber/Compound Type** |
| --- | --- | --- |
| **PH** | Psyllium Husk | Soluble, Gel-Forming Fiber |
|  | Maltodextrin | Digestible Starch |
|  | Citric Acid | Organic Acid |
|  | Natural, Artificial Flavors | Non-nutritive |
|  | Aspartame | Artificial Sweetener |
| **B3**<br>**(3-Fiber Blend)** | Stabilized Rice Bran | Insoluble Fiber and Resistant Starch |
|  | Resistant Maltodextrin | Resistant Starch Type 4 (RS4) |
|  | Resistant Potato Starch | Resistant Starch Type 2 (RS2) |
| **B5**<br>**(5-Fiber Blend)** | Stabilized Rice Bran | Insoluble Fiber and Resistant Starch |
|  | Resistant Maltodextrin | Resistant Starch Type 4 (RS4) |
|  | Resistant Potato Starch | Resistant Starch Type 2 (RS2) |
|  | Partially Hydrolyzed Guar Gum | Galactomannan |
|  | High-amylose maize resistant starch | Resistant Starch Type 2 (RS2) |
| **B5+AO**<br>**(5-Fiber + Antioxidant)** | Stabilized Rice Bran | Insoluble Fiber and Resistant Starch |
|  | Resistant Maltodextrin | Resistant Starch Type 4 (RS4) |
|  | Resistant Potato Starch | Resistant Starch Type 2 (RS2) |
|  | Partially Hydrolyzed Guar Gum | Galactomannan |
|  | High-amylose maize resistant starch | Resistant Starch Type 2 (RS2) |
|  | Green Tea Extract | Polyphenol Extract |
| **B5+Bar**<br>**(5-Fiber + Bar)** | Stabilized Rice Bran | Insoluble Fiber and Resistant Starch |
|  | Resistant Maltodextrin | Resistant Starch Type 4 (RS4) |
|  | Resistant Potato Starch | Resistant Starch Type 2 (RS2) |
|  | Partially Hydrolyzed Guar Gum | Galactomannan |
|  | High-amylose maize resistant starch | Resistant Starch Type 2 (RS2) |
|  | Date Paste | Carbohydrate |
|  | Dried Cherries | Carbohydrate |
|  | Whole Grain Oats | Whole Grain |
|  | Cashews, Walnuts | Nuts |
|  | Pumpkin Seeds, Chia Seeds | Seeds |
|  | Coconut Nectar | Simple Carbohydrate |
|  | Chocolate Liquor, Cocoa Nibs, Cocoa, Cocoa Butter | Cocoa-Derived Components |

